# Left Bundle Branch Area Pacing Preserves Mechanical Strain and Synchrony Compared to Right Ventricular Apical Pacing in an Acute Paired Preclinical Model

**DOI:** 10.1101/2025.11.27.690903

**Authors:** Emmanuel Offei, Kyoichiro Yazaki, Yuki Ishidoya, Martha Sofia Ruiz Castillo, Ava Yektaeian Vaziri, Ankur Shah, Derek J. Dosdall, Muhammad S. Khan

## Abstract

**Background:** Left bundle branch area (LBBA) pacing is emerging as a promising alternative to conventional right ventricular (RV) pacing, particularly in its ability to maintain physiological ventricular activation and enhance cardiac function. However, the effect of LBBA pacing on changes to left ventricular (LV) function in paired experiments on healthy hearts, particularly in relation to myocardial strain and time to peak systolic strain difference (TPSD), which are critical indicators of LV dysfunction, remains inadequately understood.

**Objective:** To investigate how the LV myocardial strain and TPSD change in a comparative study of LBBA pacing versus conventional RV apex pacing in the same hearts under normal physiological conditions.

**Methods:** Pre-clinical canine models (n=7) were implanted with pacing leads in the LBBA and RV apex. Functional parameters were assessed under acute pacing conditions and during sinus rhythm (SR) with echocardiographic assessment.

**Results:** Results demonstrated that LBBA pacing significantly improved global longitudinal strain (GLS: −13±2% vs. −9±2%, p = 0.0003) and global circumferential strain (GCS: −16±3% vs. −10±4%, p = 0.0119) compared to RV apical pacing. TPSD was significantly greater with RV pacing (65±10 ms vs. 24±7 ms, p = 0.0024). Compared with SR data, LBBA pacing showed no significant changes in GLS, GCS, or TPSD.

**Conclusion:** These findings suggest that LBBA pacing effectively preserves global myocardial strain and TPSD close to their values in SR, thereby contributing to enhanced overall LV function, as evidenced by improvements in ejection fraction, end-systolic volume, and end-diastolic volume.

## INTRODUCTION

Conduction system pacing has emerged as a promising alternative by recruiting the His–Purkinje network. ^1^ His bundle pacing can achieve near-normal activation but is limited by high capture thresholds and declining long-term reliability. ^2,3^ Left bundle branch area (LBBA) pacing, first described by Huang et al., ^4^ has since gained attention as a more practical and durable strategy. By directly engaging the left bundle branch, this technique offers stable capture thresholds and activation patterns that better preserve native electromechanical synchrony. ^5^ Patient studies demonstrate that LBBA pacing not only improves electrical conduction but may also better preserve left ventricular (LV) function compared with right ventricular (RV) apex pacing. ^6^ Despite recent advancements, using paired experiments on healthy hearts to study the effect of LBBA pacing on changes in LV function, particularly in relation to myocardial strain and the time to peak strain difference (TPSD)— critical indicators of LV dysfunction—remains unexplored.

Numerous clinical observational studies have examined the effect of LBBA pacing on LV function and activation; only a limited number of studies involving patients without impaired cardiac function have specifically investigated how LBBA pacing affects LV myocardial strain and mechanical synchrony in comparison to RV pacing. ^7,8^ Additionally, the existing evidence is largely observational and inadequate, particularly since myocardial strain and TPSD are often already compromised across myocardial substrates in patients before either LBBA or RV apex lead implantation. ^5,9^ To overcome this condition, we propose using a preclinical healthy canine model, free of confounding effects of pre-existing cardiac pathology, to allow a comparative study of LBBA pacing versus conventional RV apex pacing in the same hearts under normal physiological conditions. The results of our study demonstrate the preservation of LV myocardial strain and TPSD during LBBA pacing in pre-clinical healthy canine hearts compared to sinus rhythm.

## METHODS

The experimental protocol adheres to the Guide for the Care and Use of Laboratory Animals and has received approval from the Institutional Animal Care and Use Committee (IACUC) of the University of Utah.

### Animal Procedure

Seven mongrel dogs (mean weight 30 ± 2.1 kg) were pre-anesthetized with intravenous fentanyl (2–10 mcg/kg) and subsequently anesthetized with intravenous propofol (5–8 mg/kg). Anesthesia was maintained using isoflurane (1.0–3.0%) delivered via a ventilator with a mixture of 50% oxygen and 50% room air. A 12-lead ECG was placed according to standard protocol and connected to a PowerLab system (AD Instruments, Dunedin, New Zealand). Throughout the procedure, physiological parameters—such as heart rate, body temperature, blood oxygen saturation, and EtCO_₂_—were continuously monitored and maintained within normal limits. To ensure analgesia, a continuous rate infusion (CRI) of fentanyl was administered at 2–10 mcg/(kg·hr). Cefazolin (20 mg/kg, IV) was given every 90–120 minutes for prophylactic antibiotic coverage. In cases of intraoperative hypotension (mean arterial pressure <60 mmHg), a bolus of Lactated Ringer’s solution (75–120 mL) was administered intravenously. If hypotension persisted, a dopamine CRI (2–10 mcg/(kg·min)) was initiated. For sinus bradycardia, atropine (0.01–0.05 mg/kg, IV bolus) was administered to restore a normal heart rate.

### Lead and ICD Implantation

For lead implantation, the jugular vein was surgically exposed. Under fluoroscopic guidance, two pacing leads were placed: one at the right ventricular (RV) apex (5076 CapSureFix Novus MRI™, Medtronic, Inc.) and the other at the left bundle branch area (LBBA) (3830 SelectSecure™, Medtronic, Inc.) as illustrated in **Figure 1a**. Capture at the RV and LBBA sites was verified by analyzing QRS morphology changes in ECG leads V1, V5, and V6 ^10–12^ . A right bundle branch block pattern (qR) was observed in ECG lead 1, and a short stimulus to left ventricular activation time (Stim-LVAT) of 58.33 ±18.35 ms was observed in leads V5 and V6 during LBBA capture. This remained constant during low and high pacing outputs. A left bundle branch block pattern (rS) was observed in lead V1, and the Stim-LVAT was relatively long – 106.67±21.83 ms during RV pacing (**Figure 1b)**. After confirming the leads’ placement at the RV and LBBA, the leads were connected to two ICDs (Cobalt™ XT DR, Medtronic, Inc.), which were subsequently secured in a subcutaneous pocket created near the jugular access site. Pacing was conducted at twice the diastolic pacing threshold.

**Fig 1.**
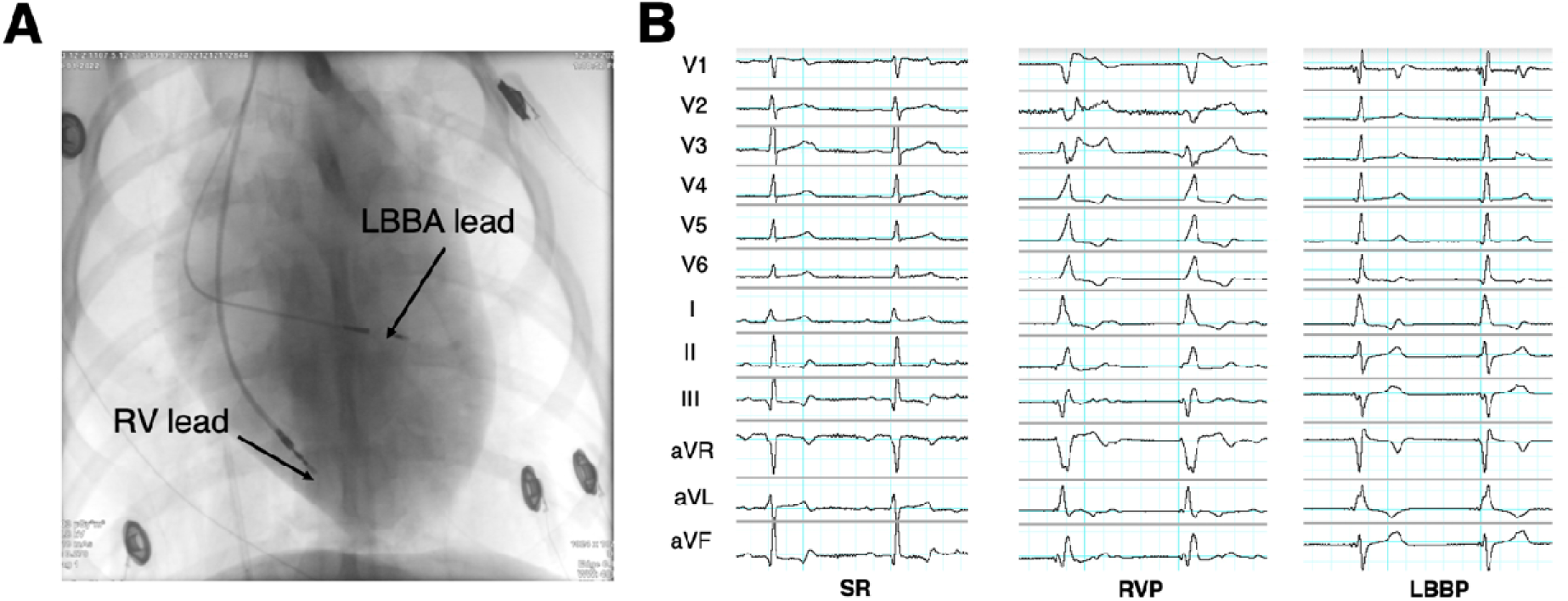
Lead implantations and ECG recordings. **(A)** Fluoroscopic projection (anterior-posterior view) showing the pacing leads implanted at the RV apex and the left bundle branch. **(B)** 12-lead ECG shows normal sinus rhythm (left), RV paced rhythm (middle), and LBB paced rhythm (right).

### Echocardiography and Electrocardiography Recordings

Following successful lead and ICD implantation, echocardiographic measurements were acquired in anesthetized dogs during SR and while pacing pulses were delivered to the RV and LBBA leads using the Siemens SC2000 Helix Ultrasound System (Siemens AG, Berlin, Germany). All measurements were collected over a minimum of 3–5 consecutive beats. After each pacing site recording, a pause of at least 30 seconds was initiated to allow for the restoration of a normal SR. Image acquisition focused on the LV short-axis view at the mid-papillary level and the apical four-chamber view. All acquired images were digitally stored for offline analysis, utilizing the built-in two-dimensional speckle tracking algorithm on the Siemens SC2000 Helix Ultrasound System, as previously described elsewhere. ^13,14^ Electrocardiography data were recorded using the 12-lead ECG during the study with the PowerLab system (AD Instruments, Dunedin, New Zealand). The standard clinical protocol, as reported elsewhere ^15^ was followed to analyze QRSd during SR, LBBA pacing, and RV pacing.

### LV Function, Myocardial Strain, and Mechanical Synchrony

Using speckle tracking echocardiography, the eSie VVI software in the Siemens SC2000 system computed various LV functional parameters: ejection fraction (EF), end-diastolic volume (EDV), end-systolic volume (ESV), global longitudinal strain (GLS), global circumferential strain (GCS), and global radial strain (GRS) from the LV apical four-chamber image. For GCS and GRS, the myocardium was traced manually from the LV short-axis images, starting at the septal region and moving circumferentially in a clockwise direction until the contour was complete. Mechanical synchrony for SR, LBBA pacing, and RV pacing was evaluated using radial strain curves from short-axis images. The time difference from the QRS onset to the peak systolic strain of the early-activated septum and the late-activated lateral wall was computed and expressed as TPSD in milliseconds (ms).

### Statistical Analysis

To identify statistically significant differences among the means of the measured variables with and without pacing strategies (SR vs. LBBA pacing vs. RV pacing) regarding LV function and mechanical synchrony, a repeated measures (RM) one-way analysis of variance (ANOVA), with the Geisser-Greenhouse correction was performed. When the ANOVA indicated a significant overall difference, a Tukey’s significant difference (HSD) post hoc test was applied for multiple pairwise comparisons between the groups. For any group that did not pass the Normality and Lognormality tests under the Kolmogorov-Smirnov test, a Kruskal-Wallis test was conducted, and Dunn’s test was then applied for multiple pairwise comparisons between the groups. Statistical significance was defined at a threshold of p < 0.05. All analyses were conducted using GraphPad Prism (GraphPad Software Inc., Boston, MA, USA).

## RESULTS

Leads were implanted at the RV apex and the LBBA in all seven canines enrolled in the study (**Figure 1a**). One animal was excluded due to the poor quality of the recorded echocardiographic images. QRS duration (QRSd) was not measured in one of the animals due to significant noise in the ECG leads.

When paced at the LBBA, LVEF did not significantly differ from that during SR (34±9 vs. 42±7 %, *p* = 0.1194), as shown in **Figure 2a**. However, LVEF during RV (30±8%) pacing was significantly lower compared to SR (42±7%, *p* = 0.0126) and LBBA pacing (34±9%, *p* = 0.0073). Similarly, LVESV did not show statistical differences between SR and LBBA pacing (30±9 vs. 33±6 mL, *p* = 0.5131, **Figure 2b**). An elevated LVEDV in **Figure 2c** indicates an overloaded heart, common in heart failure conditions. This difference was not significant between SR and LBBA pacing (51±10 vs. 53±8 mL, *p* = 0.4179). In contrast, RV pacing resulted in a significantly higher LVESV (40±7 mL) and LVEDV (56±7 mL) compared to SR (LVESV: 30±9 mL, *p* = 0.0206 and LVEDV: 51±10 mL, *p* = 0.0285) and LBBA pacing (LVESV: 33±6 mL, *p* = 0.0314 and LVEDV: 53±8 mL, *p* = 0.0076), as shown in **Figure 2b** and **2c**, respectively.

**Fig. 2.**
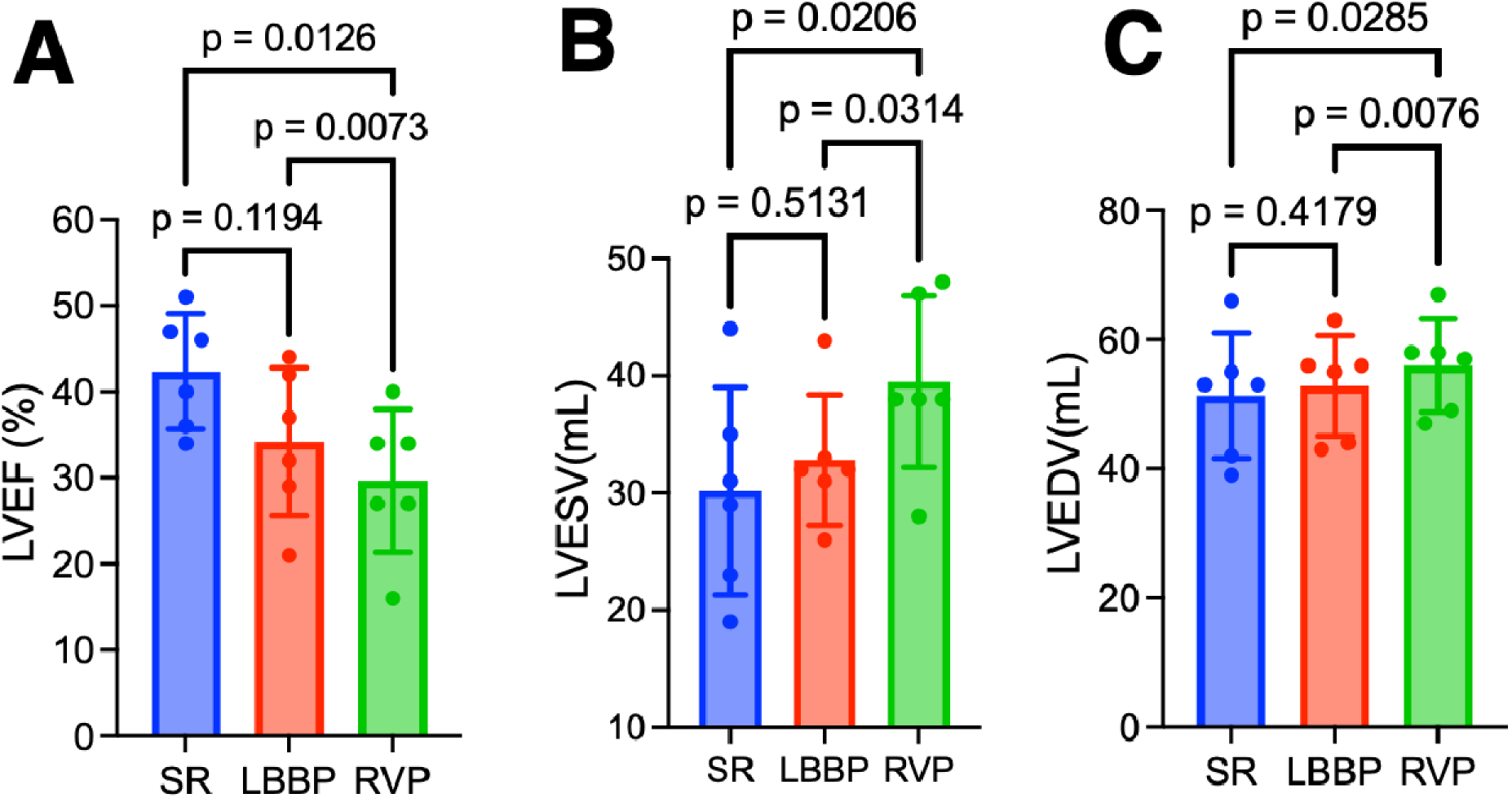
LV function comparison between SR, LBBP, and RVP (n = 6). **(A)** Left ventricular ejection fraction (LVEF). **(B)** Left ventricular end systolic volume (LVESV). **(C)** Left ventricular end diastolic volume (LVEDV). Results were quantified from the 4-chamber apical view of the LV. Statistical comparison was performed using a RM one-way ANOVA with the Geisser-Greenhouse correction followed by post hoc Tukey’s multiple comparison tests to compare between groups. Statistical comparison was performed using a RM one-way ANOVA with the Geisser-Greenhouse correction followed by post hoc Tukey’s multiple comparison tests to compare between groups. Results are presented as mean±SD.

In our study of healthy canine LV myocardium, the GLS outcome for SR and LBBA pacing was not statistically different (−16±3 vs. −13±2 %, *p* = 0.0607, **Figure 3a**). In contrast, GLS during RV pacing was significantly lower than both SR (−9±2% vs. −16±3 %, *p* = 0.0015) and LBBA pacing (−9±2% vs. −13±2%, *p* = 0.0003). Similarly, GCS was not significantly different for SR compared to LBBA pacing (−19±6% vs. −15±4 %, *p* = 0.0747, **Figure 3b**). However, the GCS during RV pacing (−10±4%) was considerably lower compared to SR (−19±6 %, *p* = 0.0109) and LBBA pacing (−15±4 %, *p* = 0.0119). For GRS data, the results showed a different trend as shown in **Figure 3c**. Compared to SR (41±8%), GRS was significantly lower for LBBA pacing (24±2 %, *p* = 0.0095) and RV pacing (18±7 %, *p* = 0.0070). It is noted that GRS was relatively higher for LBBA pacing compared to RV pacing, although this difference did not achieve statistical significance (*p* = 0.1321).

**Fig. 3.**
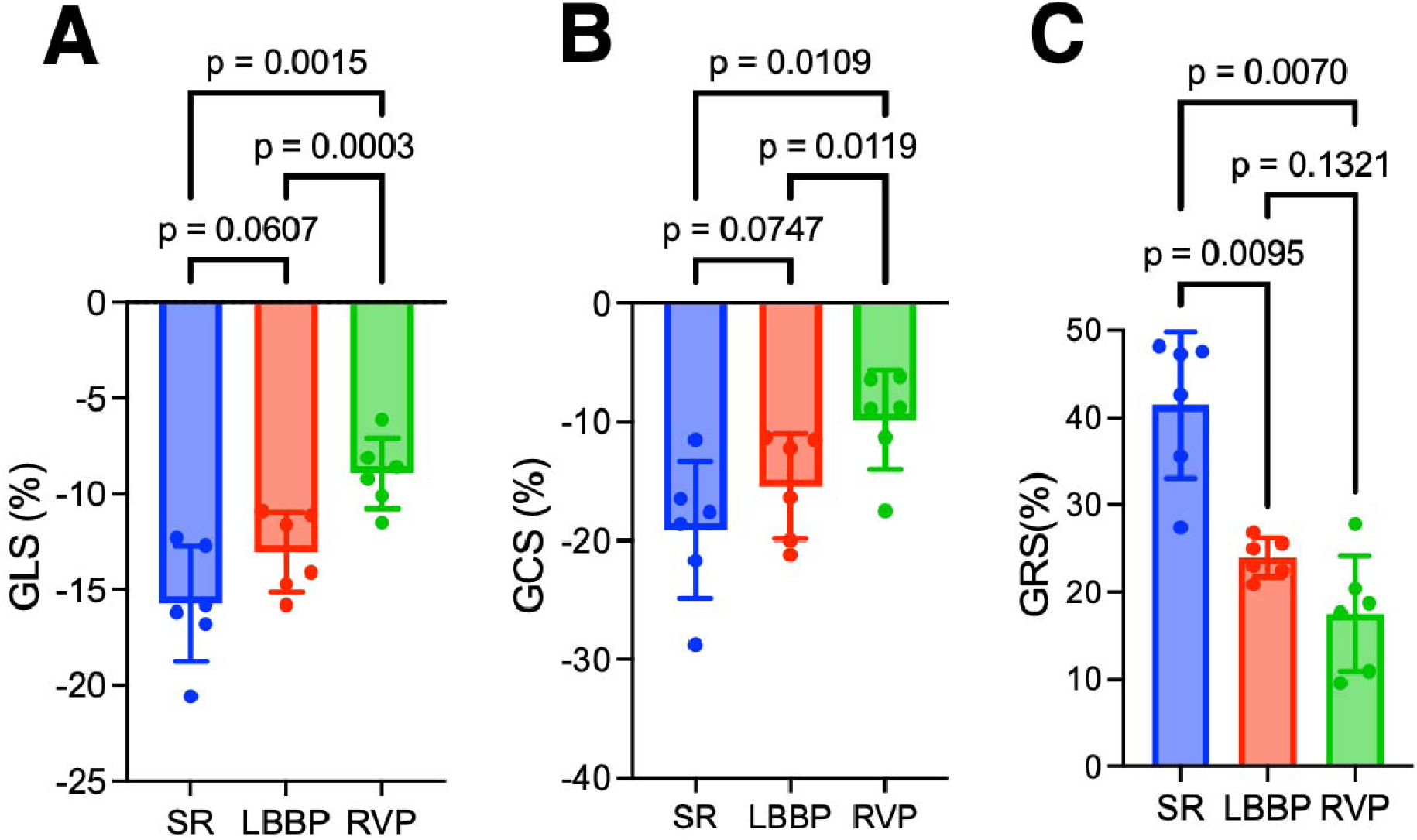
Global myocardial strains from the 4-chamber apical and short-axis (papillary level) views. **(A)** Global longitudinal strain (GLS), **(B)** Global circumferential strain (GCS), **(C)** Global radial strain (GRS). Statistical comparison was performed using a RM one-way ANOVA with the Geisser-Greenhouse correction followed by post hoc Tukey’s multiple comparison tests to compare between groups. Results are presented as mean±SD.

The TPSD within the LV measures the timing difference in mechanical contraction between the early-activated septum and the late-activated lateral wall (**Figure 4a**), serving as a measure for LV mechanical synchrony. We hypothesized that in healthy myocardium, the LBBA pacing promotes synchronous activation of the LV and shortens the timing between septal and lateral wall contraction. The TPSD for SR was not significantly different from that for LBBA pacing (21±10 vs. 24±7 ms, *p* = 0.1260), as shown in **Figure 4b**. In contrast, TPSD was significantly longer for RV pacing (65±10 ms) compared to SR (21±10 ms, *p* = 0.0038) and LBBA pacing (24±7 ms, *p* = 0.0024).

**Fig. 4.**
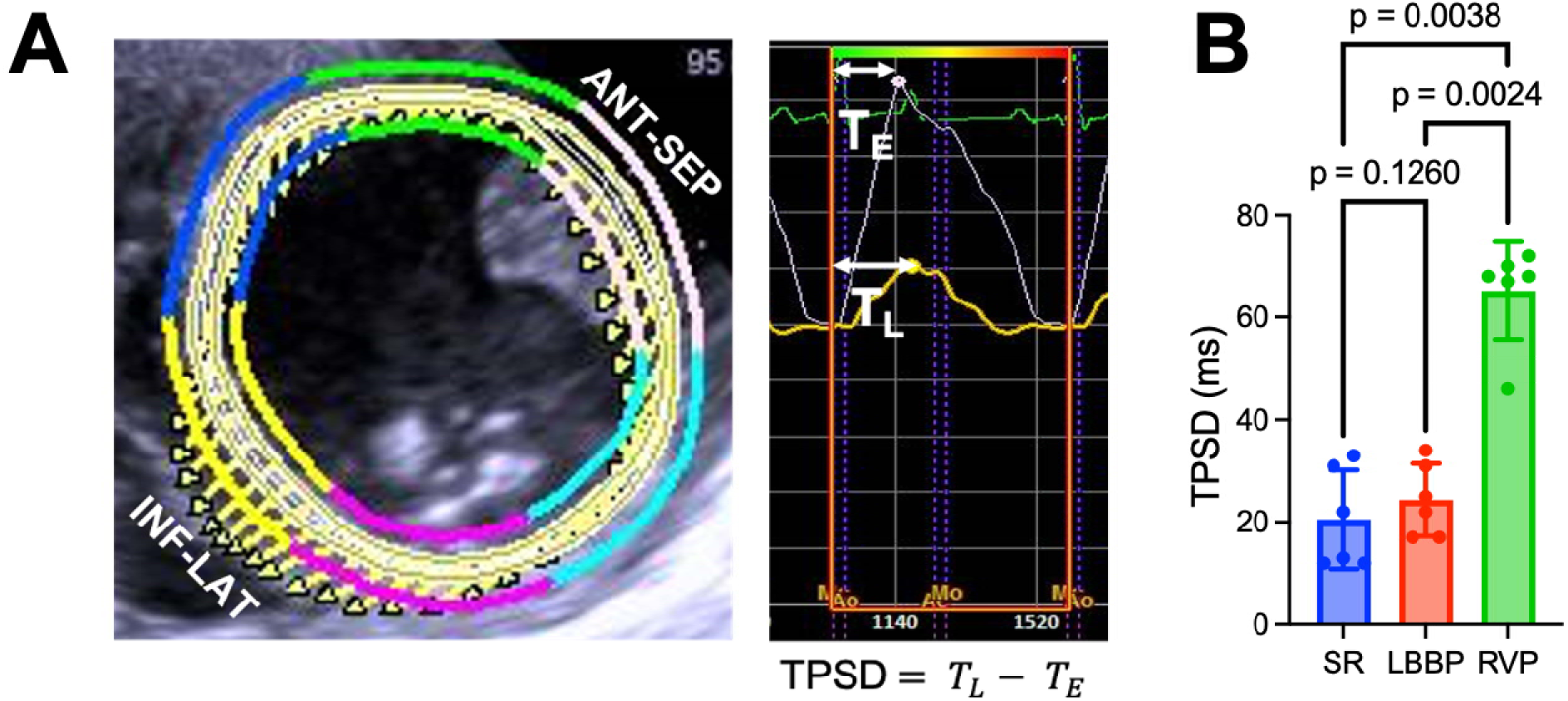
LV mechanical dyssynchrony, also known as time to peak strain difference (TPSD), was calculated using peak strain time difference. **(A)** Representative image from a short-axis view of the LV at the papillary level showing the anteroseptal (ANT-SEP) and inferolateral (INF-LAT) segments. **(B)** Peak radial strain is used to measure mid-level short-axis time-to-peak difference between anteroseptal (ANT-SEP) and inferior-lateral (INF-LAT) segments. T*_L_* and T*_E_* represent time-to-peak strain ‘late’ and ‘early’, respectively. **(C)** A Kruskal-Wallis test was conducted to test for a difference between groups (n = 6). Post hoc Dunn’s multiple comparison tests results are shown with their p-values. Results are presented as mean±SD.

A prolonged QRS duration may indicate conduction abnormalities, which are often associated with ventricular remodeling, particularly in conditions such as heart failure or bundle branch block. Pacing from the LBBA led to a narrowing of the QRS complex, as shown in **Figure 5**; however, this change was not significantly different from the QRSd during SR (71±8 vs. 58±18 ms, *p* = 0.4561). In contrast, when pacing from the RV apex (111±11 ms), the QRSd significantly increased compared to SR (58±18 ms, *p* = 0.0164) and during LBBA pacing (71±8 ms*, p* = 0.0161).

**Fig. 5.**
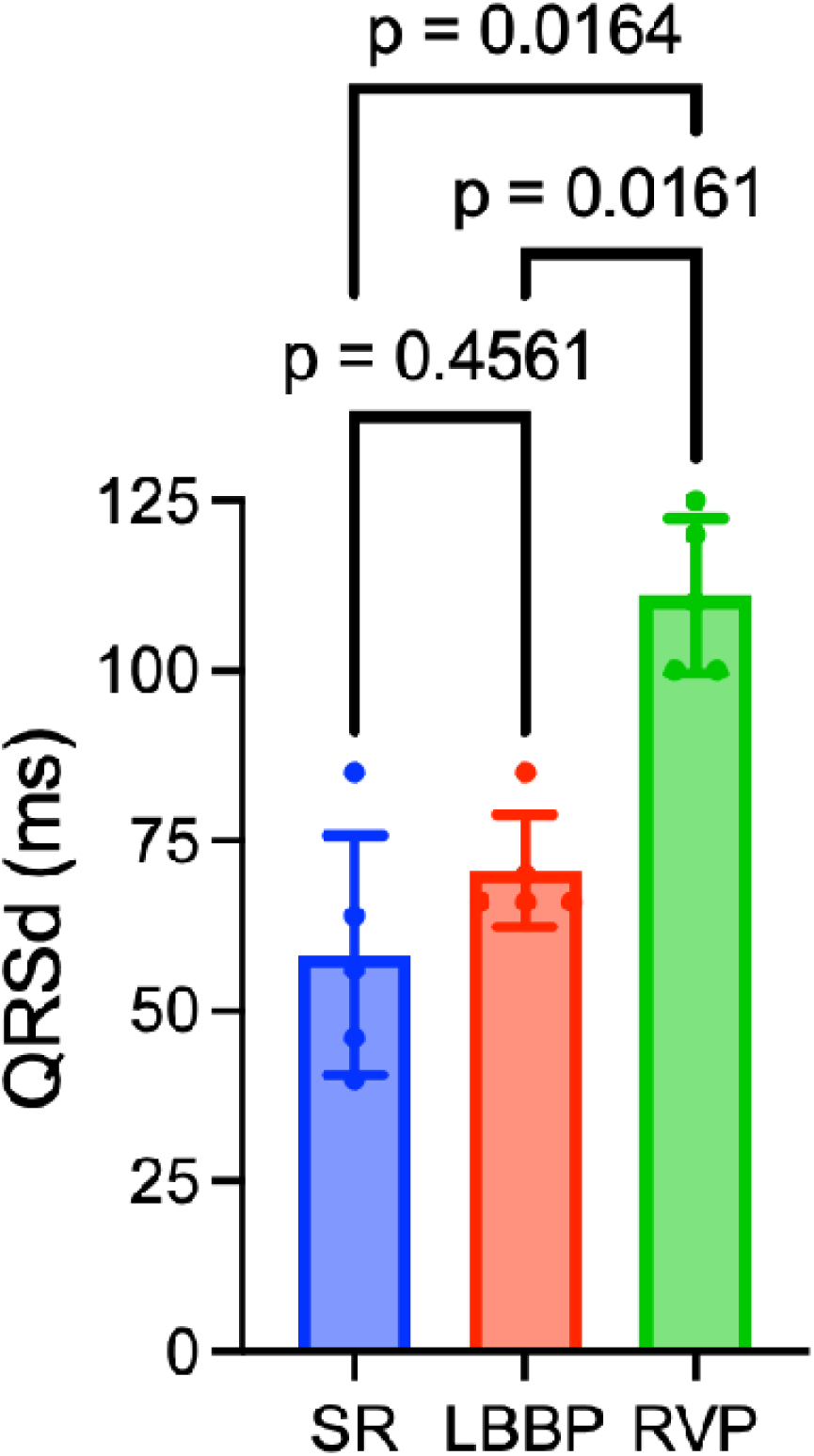
QRS duration (QRSd) in sinus and during pacing (n = 5) was quantified using a 12-lead ECG. Statistical comparison was performed using a RM one-way ANOVA with the Geisser-Greenhouse correction followed by post hoc Tukey’s multiple comparison tests to compare between groups. Results are presented as mean±SD.

## DISCUSSION

Our study provides a direct comparison of the heart’s electrical and mechanical function for RV pacing versus LBBA pacing, utilizing data from the same healthy animals. Heterogeneity in the baseline characteristics of patient data confounds direct comparison of pacing location, while simultaneous dual lead implantation in uncompromised hearts offers a direct comparison of the therapeutic effect of LBBA versus RV pacing. Our study demonstrated similar LV myocardial strain and TPSD during LBBA pacing compared to sinus rhythm, and improved measures compared to RV pacing. Synchronous ventricular activation, achieved through LBBA pacing, is more likely to maintain LVEF close to that during SR ^16,17^; however, variability between patient groups due to differing heart conditions hinders the understanding of the physiological impact of LBBA pacing compared to RV pacing on LV myocardial strain and TPSD.

Previous clinical studies demonstrated that LBBA pacing delivers electrical activation via the native conduction system, thereby promoting synchronous LV activation and preserving cardiac function. ^18,19^ Our results extend these observations and support that preferential recruitment of myocardial fibers in the longitudinal and circumferential directions during LBBA pacing improves LV mechanical synchrony and maintains the systolic function. We hypothesized that the activation spreading throughout the conduction system would facilitate mechanical synchrony between the LV septal and lateral walls, leading to improved LV -GLS and -GCS, a phenomenon that was not observed during RV pacing. LBBA pacing, therefore, preserves LV mechanical function better than RV apical pacing. To our knowledge, this is the first experimental demonstration of strain-dependent preservation of LV function in healthy canine hearts, highlighting the mechanistic advantage of LBBA pacing over RV pacing. LBBA pacing preserved LV systolic function, whereas RV pacing impaired systolic performance, evidenced by reduced LVEF (**Figure 2a**) and increased LVEDV (**Figure 2b**) and LVESV (**Figure 2c**). These findings are consistent with the notion that LBBA pacing recruits the LV through rapid conduction system activation rather than slow cell-to-cell propagation, resulting in synchronous contraction. ^20^ Multiple investigations showed that LBBA pacing preserves or improves LVEF, including reports by Sousa et al., ^21^ Su et al., ^22^ and Vijayaraman et al. ^16^ Similarly, Ye et al. observed a reduction in LVESV of 12 mL at 6-month follow-up in atrial fibrillation patients paced at the LBBA. ^23^ In contrast, RV pacing disrupts synchrony, reducing contractility and leading to pacing-induced cardiomyopathy in a subset of patients. ^24–27^ Although these clinical data are highly supportive, demonstrating that LBBA pacing prevents maladaptive LV remodeling, exploring the physiological effects of LBBA pacing compared to RV pacing on myocardial strain and TPSD in patients with impaired hearts may be challenging. Our findings indicate that myocardial strain data effectively identify subtle or early changes in myocardial contractility associated with LBBA pacing. For instance, both GLS (**Figure 3a**) and GCS (**Figure 3b**) were significantly reduced during RV apical pacing compared with SR and LBBA pacing, which may indicate impaired longitudinal and circumferential mechanics in healthy myocardium. Conversely, LBBA pacing preserved GLS and GCS close to SR baseline values, reflecting synchronous myocardial fiber recruitment. Our observations agree with prior reports showing deterioration of GLS with RV apical pacing ^28,29^ and greater preservation of strain with LBBA pacing. ^30^ Similarly, Mao et al. also noted a greater decline in GLS after one year of RV pacing than LBBA pacing (−4.8% vs. −1.4%) in symptomatic bradycardia patients. ^31^ Nevertheless, GRS (**Figure 3c**) was significantly reduced during LBBA and RV pacing compared with SR, though not significantly different between the pacing modalities (LBBA vs. RV). These results suggest that radial thickening is particularly sensitive to non-physiological activation and may require further mechanistic investigation. Similarly, Yao et al. reported no significant GRS differences between RV septal pacing and LBBA pacing, despite improvements in their data acquired for GLS and GCS. ^30^ Additionally, with improved GLS and GCS, LBBA pacing preserves the physiologic timing of septal and lateral wall contraction as indicated in our TPSD data (**Figure 4b**), a marker of LV synchrony. Clinical studies by Sun et al. ^9^ and Yang et al. ^5^ similarly reported shorter time-to-peak strain intervals with LBBA pacing compared to RV pacing, highlighting its superior ability to maintain mechanical synchrony. Besides the LV mechanics advantage, we also observed that QRSd (**Figure 5**) remained narrow with LBBA pacing, in contrast to QRS widening with RV pacing. This reflects rapid impulse propagation via the His-Purkinje system, which conducts at ∼2-3 m/s compared to 0.5-1.0 m/s in working myocardium. ^32,33^ Prior studies confirm these findings, showing narrower QRS complexes during LBBA pacing compared to RV pacing. ^5,34^

## CONCLUSION

The findings of the study underscore the advantages of LBBA pacing as it significantly facilitates synchronous activation of the LV and enhances myocardial contraction. Compared to RV pacing, by recruiting the longitudinal and circumferential fibers throughout the LV and consequently reduces time-to-peak strain intervals. LBBA pacing not only modulates global LV myocardial strain positively but also promotes LV systolic performance. These results provide compelling evidence for the use of LBBA pacing as a superior therapeutic approach for patients with impaired LV function, suggesting it may enhance overall cardiac performance and patient outcomes.

## LIMITATIONS

Our study was limited by a small sample size and was primarily focused on the acute effects of pacing the LBBA and the RV apex on LV function. We did not investigate the long-term effects or clinical benefits of pacing both sites, as our main objective was to examine myocardial strain and TPSD in the paired healthy canine models. Data collection occurred while the subjects were anesthetized, which may have led to lower cardiac function values due to the effects of isoflurane ^35,36^. Nonetheless, our conclusions remain robust, as the findings were consistent across different operators, reproducible in repeated analyses, and corroborated by QRS duration data.

## Acknowledgement

The authors would like to thank Orvelin Roman for performing surgical procedures related to this work.

## Authorship

All authors attest they meet the current ICMJE criteria for authorship.

## Ethics Statement

The experimental protocol adhered to the ARRIVE guidelines and the Guide for the Care and Use of Laboratory Animals and was approved by the Institutional Animal Care and Use Committee of the University of Utah.

